# Comparative genomics-driven design of virus-delivered short RNA inserts triggering robust gene silencing

**DOI:** 10.1101/2025.06.03.657697

**Authors:** Arcadio García, Verónica Aragonés, Silvia Gioiosa, Francisco J. Herraiz, Paloma Ortiz-García, Jaime Prohens, José-Antonio Daròs, Fabio Pasin

## Abstract

Engineered RNA viruses can downregulate target genes through virus-induced gene silencing (VIGS). Largely optimized using the model plant *Nicotiana benthamiana*, current VIGS protocols rely on viral vectors engineered to deliver 200-400 nt inserts with homology to a target gene, often located in less conserved regions to ensure specificity. Given the *N. benthamiana* allopolyploidy and functional redundancy, we leveraged enhanced genomics and transcriptomics resources to obtain curated locus annotations, which guided the design of virus-delivered short RNA inserts (vsRNAi) for simultaneous targeting of homeologous gene pairs. Our results showed that quantitative silencing phenotypes could be obtained using 24-, 28-, and 32-nt vsRNAi that targeted conserved regions of functionally redundant gene pairs. vsRNAi triggered informative transcriptome-wide changes, and target transcript downregulation was linked to gene-pair-specific production of 21- and 22-nt sRNAs. The portability of vsRNAi to crops was confirmed in tomato (*Solanum lycopersicum*) and scarlet eggplant (*Solanum aethiopicum*). Overall, our results support the usefulness of vsRNAi-based approaches in functional genomics and trait reprogramming of the model species N. benthamiana, as well as of crops that contribute to global and local food provisions.

**HIGHLIGHTS**

- Comparative genomics enables design of virus-delivered short RNA inserts (vsRNAi)
- 32-, 28-, and 24-nt vsRNAi resulted in quantitative gene silencing phenotypes
- vsRNAi trigger transcriptome-wide changes for gene functional characterization
- vsRNAi elicit production of 21- and 22-nt small RNAs and gene downregulation
- vsRNAi approaches assist functional genomics and phenotypic alteration in crops

## MAIN TEXT

RNA products and RNA-virus based technologies have the potential to transform agriculture by enabling on-demand crop trait reprogramming and effective pest and disease management (Pasin *et al*., 2024; Rössner *et al*., 2022). In virus-induced gene silencing (VIGS), engineered RNA viruses can redirect the host RNA interference machineries to target gene silencing through the production of gene-specific small RNAs (sRNAs) (Rössner *et al*., 2022). Although endogenous sRNAs and those resulting from VIGS are in the 20–30-nt range, VIGS vectors are engineered to deliver larger inserts of 200–400 nt with homology to a target gene, often located in less conserved regions to ensure specificity (Ahmed *et al*., 2020). Reducing insert sizes, which are currently nearly 10 times larger than end products required for RNA interference, may overcome the need for biological material for insert amplification, and be amenable for low-cost DNA chemical synthesis, thereby enhancing the VIGS scalability and applicability to non-model species.

*Nicotiana benthamiana* is the most widely used model for optimizing VIGS protocols. However, this host has a complex, allotetraploid genome with functional redundant homeologous gene pairs, and for which no high-quality assemblies were available until very recently (Ranawaka *et al*., 2023). Multi-gene CRISPR-Cas9 mutagenesis was applied to tackle functional redundancy in plants (Berman *et al*., 2025; Ellison *et al*., 2020). Here, we hypothesized that VIGS insert sizes could be lowered to match those of endogenous sRNAs by combining enhanced genomics and transcriptomics resources to guide the design of virus-delivered short RNA inserts (vsRNAi) for simultaneous targeting of homeologous gene pairs.

We focused on the magnesium protoporphyrin chelatase subunit I (*CHLI*) gene, whose downregulation results in leaf yellowing due to a reduction of chlorophyll biosynthesis and levels. Tomato (*Solanum lycopersicum*) is a diploid relative of *N. benthamiana* with high-quality genome assembly and annotation. The tomato *CHLI* coding sequence (CDS) is distributed across three exons with boundaries supported by transcriptomic expression analysis (Supporting experimental procedures). Analysis of reported *N. benthamiana* Niben101 and Niben261 annotations however revealed *CHLI* variation including large sequence deletion and insertion, not supported by results obtained by *de novo* transcriptome assembly (Figure S1). A high-quality chromosome-level genome assembly was recently reported for *N. benthamiana* (Ranawaka *et al*., 2023). Sequence searches using the tomato CHLI protein identified homolog loci in the chromosomes 5 (NbLab360C05; evalue = 2.13×10^−210^) and 10 (NbLab360C10; evalue = 1.26×10^−211^), whose expression and gene structures were validated by mapping sequencing reads from rRNA-depleted total RNA or polyA-tailed RNA samples. Leveraging this curated annotation, vsRNAi were designed to target CDS regions conserved in *N. benthamiana* and tomato *CHLI* (Figure S1).

Custom-synthesized DNA oligonucleotide pairs spanning vsRNAi sequences were inserted into pLX-TRV2 of the JoinTRV vector system, which is based on tobacco rattle virus (TRV) (Aragonés *et al*., 2022), by one-step digestion-ligation reactions (Figure 1a). JoinTRV derivatives with 32-, 28-, 24- and 20-nt inserts targeting *CHLI* (vCHLI, vCHLI-28, vCHLI-24 and vCHLI-20, respectively) were inoculated to *N. benthamiana*. After 10 days, upper uninoculated leaves of plants treated with vCHLI, vCHLI-28, and vCHLI-24 showed a yellowing phenotype. Fluorometry revealed a significant reduction of chlorophyll levels in vCHLI 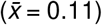, vCHLI-28 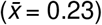, and vCHLI-24 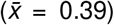 compared to controls (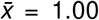; Figure 1a), and a significant positive correlation with *CHLI* transcript levels measured by RT-qPCR (Pearson’s *r* = 0.892; *p* = 0.001; Figure S2).

**Figure 1.**
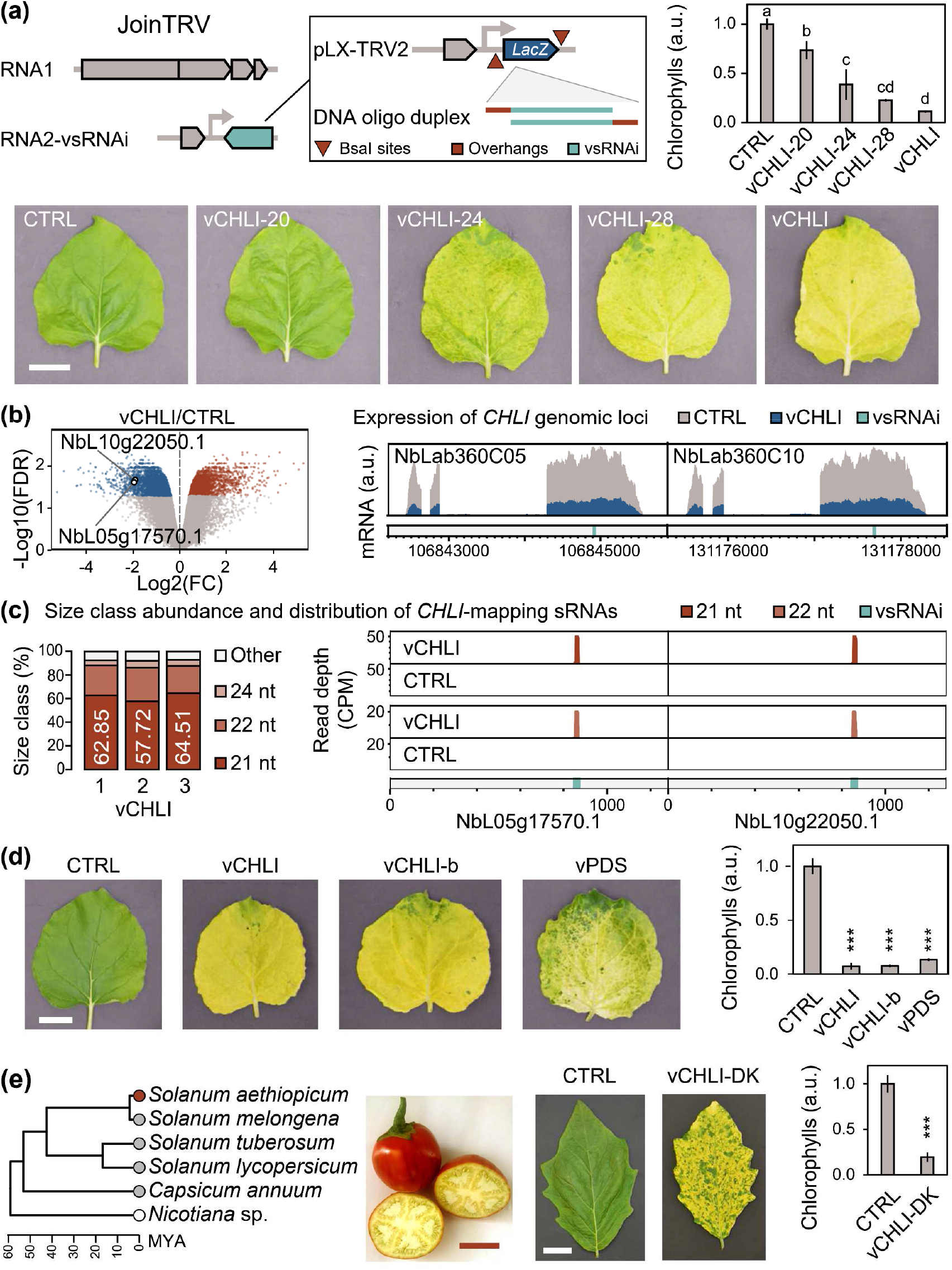
Virus-delivered short RNA inserts (vsRNAi) trigger gene silencing. **(a)** JoinTRV vector assembly and delivery of 32-, 28-, 24- and 20-nt vsRNAi targeting the two *N. benthamiana CHLI* homeologues (vCHLI, vCHLI-28, vCHLI-24, and vCHLI-20). Leaf phenotypes and chlorophyll levels (mean ± SD, *n* = 3) are shown; CTRL, control. Different letters indicate significant differences (*p* < 0.05), by one-way ANOVA and Tukey’s honestly significant difference (HSD) test. **(b)** Transcriptomics of vCHLI and CTRL samples (*n* = 3) detects a significant downregulation of *CHLI* expression (FDR < 0.05). **(c)** In vCHLI samples, small RNA sequencing shows predominance of 21- and 22-nt sRNAs mapping to *CHLI*, and with a read depth peak (mean, *n* = 3) localized to the vsRNAi-targeted region. **(d)** Leaf phenotypes and chlorophyll levels (mean ± SD, *n* = 3) are shown for vCHLI-b and vPDS, targeting *CHLI* and *PDS* homeologues, respectively; ∗∗∗, *p* < 0.001, Student’s *t*-test. **(e)** vsRNAi portability to scarlet eggplant (*Solanum aethiopicum*). Phylogeny and fruits are shown, alongside leaf phenotypes and chlorophyll levels (mean ± SD, *n* = 3) of “Rossa di Rotonda” plants treated with vCHLI-DK, a pTRV1 + pTRV2 derivative targeting *CHLI*; ∗∗∗, *p* < 0.001, Student’s *t*-test.

Silencing phenotypes of vCHLI-treated plants were robust and equivalent to those obtained by using a VIGS vector including a 300-nt *CHLI* cDNA fragment (Figure S3). Transcriptomes of plants inoculated with the unmodified JoinTRV (CTRL) or its vCHLI derivative were analyzed. Abundance of over 4,000 transcripts was significantly altered in the vCHLI/CTRL comparison (FDR < 0.05; Figures 1b, S4; Data S1, S2). Transcriptome-wide functional analysis revealed an enrichment of gene ontology terms associated to light responses (GO:0009416, *p* = 4.60×10^−12^), and biological processes involved in carbohydrate metabolism (GO:0005975, *p* = 0.0023), cellulose biosynthesis and cell wall biogenesis (GO:0030244, *p* = 4.80×10^−10^; GO:0042546, *p* = 6.10×10^−09^) (Figure S4c; Data S3-S5), consistent with the anticipated reduction of cell photosynthetic capacity caused by the *CHLI* downregulation.

Among downregulated transcripts, we identified the *CHLI* homeologues (NbL05g17570.1, log2(FC) = −1.97, FDR = 0.024; NbL10g22050.1, log2(FC) = −1.92, FDR = 0.021), and confirmed a global expression reduction of their genomic loci beyond the 32-nt region targeted by vCHLI (Figure 1b; Data S6). Viral amplification of large VIGS inserts can lead to overestimation of expression levels of homologous host genes (Figure S5, S6); in contrast, vsRNAi enable transcriptome-wide quantification of target gene silencing.

We next sequenced sRNAs to assess if their production is involved in the vCHLI-induced phenotypic and transcriptomic changes. In vCHLI samples, vsRNAi triggered host-derived production of sRNAs (Figure S7), and a marked enrichment of sRNAs mapping to *CHLI* transcripts (NbL05g17570.1, log2(FC) = 7.98, *p* = 0.020; NbL10g22050.1, log2(FC) = 9.23, *p* = 0.020) (Figure S8; Data S7, S8). 21-nt sRNAs were predominant (57.72%-64.51% range; *n* = 3), followed by 22-nt sRNAs (23.24%-28.46% range) (Figure 1c; Data S9), and their levels showed a significant negative correlation with those of *CHLI* transcripts (Pearson’s *r* ≤ −0.888; *p* ≤ 0.021; Figure S9). The vsRNAi-targeted region of *CHLI* transcript sequences exhibited a localized accumulation of 21- and 22-nt sRNAs in vCHLI samples, which was absent in control samples. (Figure 1c; Data S10). We concluded that vsRNAi trigger target gene downregulation through region-specific enrichment of 21- and 22-nt sRNAs, known end products of Dicer-like 4 (DCL4) and DCL2, respectively (Rössner *et al*., 2022).

We used vCHLI-b and vPDS to target a second 32-nt region conserved in *CHLI* homeologues or 32-nt of the *PHYTOENE DESATURASE* (*PDS*) gene pair, respectively. In the upper uninoculated leaves of treated plants, we observed a reduction in green pigmentation and chlorophyll levels (Figure 1d), confirming the broad applicability of vsRNAi for *N. benthamiana* gene functional characterization.

We next assessed vsRNAi portability to crops. The vCHLI insert is conserved in tomato *CHLI* (Figure S1); “Moneymaker” seedlings inoculated with vCHLI showed leaf yellowing and a significant reduction of chlorophyll levels compared to control plants (Figure S10a). Scarlet eggplant (*Solanum aethiopicum*; Figure 1e) is an underutilized solanaceous crop mostly grown in Africa and Brazil with limited availability of genetic resources and biotechnological tools (Gramazio *et al*., 2016). Searches of *de novo* assembled transcriptomes and of a recently released genomic assembly (Benoit *et al*., 2025) confirmed that the 32-nt insert of vCHLI is conserved in scarlet eggplant *CHLI* (32/32, 100% identity) (Figure S10b). Seedlings of the ecotype “Rossa di Rotonda” inoculated with vCHLI showed leaf yellowing and a significant reduction of chlorophyll levels compared to control plants (Figure S10b). Using vCHLI-DK, a derivative of the pTRV1 plus pTRV2 vector system used for VIGS and genome editing (Ellison *et al*., 2020), we delivered a *CHLI*-targeting 32-nt vsRNAi and obtained equivalent results (Figures 1e, S10c), highlighting the versatility of our approach.

Overall, our results demonstrate that vsRNAi as short as 24 nt can effectively produce phenotypic alterations. Use of 32-nt vsRNAi results in robust gene silencing phenotypes, informative transcriptome-wide changes, and target transcript downregulation linked to gene-specific production of 21- and 22-nt sRNAs.

Viral delivery of artificial micro RNAs or trans-acting small interfering RNAs reduces VIGS insert sizes and off-targets (Cisneros *et al*., 2025). vsRNAi offer equivalent specificity and greatly simplify viral vector engineering by eliminating intermediate cloning steps and precursor elements required to activate sRNA production.

Simplified cloning of vsRNAi fragments, which are nearly 10-fold smaller than those of conventional VIGS and can be synthesized at low cost, may enable high-throughput functional genomics in model plants and crops that contribute to global and local food security. Finally, innovations implementing vsRNAi designs may provide cost-effective solutions to improve crop performance and sustainable production by on-demand alteration of crop traits, and selective control of crop pests and diseases.

## AUTHOR CONTRIBUTIONS

F.P. conceived the study; J.-A.D. and F.P. obtained funding and computational resources; A.G. and F.P. designed experiments; A.G., V.A., P.O.-G. and F.P. performed the experiments; F.J.H. and J.P. provided materials; A.G., S.G. and F.P. analyzed and managed data. F.P. wrote the manuscript with input from A.G.; all authors revised and approved the final version.

## ACKNOWLEDGEMENTS

This work was supported by grants PID2023-146418OB-I00 from Ministerio de Ciencia, Innovación y Universidades (Spain) through the Agencia Estatal de Investigación, and PROMETEO CIPROM/2022/21 from Generalitat Valenciana. A.G. is the recipient of a predoctoral contract (FPU20/05477) from Ministerio de Ciencia, Innovación y Universidades (Spain). F.P. is financed by the ‘Ramón y Cajal’ program (RYC2023-045411-I) of Ministerio de Ciencia, Innovación y Universidades (Spain), and gratefully acknowledges the grants MiniVi (ELIXIR-IIB, Cineca, Italy) and BCV-2023-1-0021 (Red Española de Supercomputación, Spain), and resources provided by Centro de Supercomputación de Galicia (CESGA, Spain). We thank Cristina Mallor Giménez (BGHZ-CITA, Spain) for *S. aethiopicum* seeds.

## DATA AVAILABILITY STATEMENT

Supplemental information accompanies this article. The transcriptomic and small RNA sequencing datasets generated are deposited under NCBI BioProject PRJNA1217923. Sequences of vsRNAi-targeted transcripts and assembled viral vectors are in Data S11 and Data S12, respectively; pLX-TRV2-vCHLI is available at Addgene (239842, https://www.addgene.org/239842/).

## SUPPLEMENTAL INFORMATION

Additional supporting information may be found online in the Supporting Information section at the end of the article.

### Appendix S1 file, including

### Supporting experimental procedures

**Table S1**. Transcriptomics and genomics resources used in the study.

**Table S2**. Software and algorithms used in the study.

**Table S3**. Oligonucleotides used in the study.

**Table S4**. Oligonucleotide duplexes used for vsRNAi vector assembly.

**Table S5**. Summary of the transcriptome sequencing (RNA-seq) datasets generated.

**Table S6**. Summary of the small RNA sequencing (sRNA-seq) datasets generated.

**Figure S1**. Annotation of *Nicotiana benthamiana CHLI* genomic loci and design of vsRNAi.

**Figure S2**. RT-qPCR of *N. benthamiana CHLI* transcripts.

**Figure S3**. Delivery of a 32-nt vsRNAi triggers robust silencing phenotypes.

**Figure S4**. Functional analysis of transcriptome-wide changes triggered by vsRNAi.

**Figure S5**. vsRNAi enable transcriptome-wide quantification of target gene silencing.

**Figure S6**. Viral amplification of large VIGS inserts interferes with quantification of target genes.

**Figure S7**. vsRNAi trigger host-derived production of sRNAs.

**Figure S8**. Production of gene-specific sRNAs triggered by vsRNAi.

**Figure S9**. Abundance of *CHLI*-mapping sRNAs negatively correlated with *CHLI* expression.

**Figure S10**. Portability of vsRNAi approaches to crops and viral vector systems.

**Supporting references**

### Supporting data file, including

**Data S1**. Transcript quantification results obtained by Salmon analysis of RNA-seq datasets from CTRL and vCHLI samples; transcript per million (TPM) values are show.

**Data S2**. Transcript differential expression values obtained by edgeR (vCHLI versus CTRL), and matched *Arabidopsis thaliana* proteins by DIAMOND blastx.

**Data S3**. GO term enrichment significance associated to transcriptome-wide changes induced by vCHLI versus CTRL.

**Data S4**. GO term enrichment significance associated to genes downregulated by vCHLI versus CTRL.

**Data S5**. GO term enrichment significance associated to genes upregulated by vCHLI versus CTRL.

**Data S6**. Expression of *CHLI* genomic loci in vCHLI and control samples; average values of normalized RNA-seq read depths are shown (*n* = 3).

**Data S7**. Transcript sequencing depth by mapping sRNA-seq reads of CTRL and vCHLI samples; counts per million (CPM) values are shown.

**Data S8**. *CHLI* sequencing depth by mapping sRNA-seq reads of CTRL and vCHLI samples; counts per million (CPM) values are shown.

**Data S9**. Size class abundance of *CHLI*-mapping sRNA reads of vCHLI samples.

**Data S10**. Distribution of 21- and 22-nt sRNAs mapped to the coding sequences of NbL05g17570.1 and NbL10g22050.1; average CPM values (*n* = 3) of vCHLI and CTRL samples are shown.

**Data S11**. Sequences of vsRNAi-targeted transcripts.

**Data S12**. Sequences of the vsRNAi vectors used in the study.

