## Appendix S1 for "Comparative genomics-driven design of virus-delivered short RNA inserts triggering robust gene silencing"

### CONTENTS

**Supporting experimental procedures**
**Supporting Tables S1-S6**
**Supporting Figures S1-S10**
**Supporting references**

### SUPPORTING EXPERIMENTAL PROCEDURES

#### Plants

Plants of *Nicotiana benthamiana*, *Solanum lycopersicum* cv. “Moneymaker” and *Solanum aethiopicum* ecotype “Rossa di Rotonda” were grown in soil mixture (3 parts of potting substrate and 1 part of vermiculite) and maintained in a greenhouse chamber at ≈24°C under a 16-h-day/8-h-night photoperiod.

#### Computational resources

Bioinformatics analyses were done in high performance computing systems through a command-line interface (Castrignanò *et al.*, 2020). Transcriptomics and genomics datasets, and software/algorithms used in the study are listed in Tables S1 and S2.

#### Comparative genomics and gene annotation

Genome sequences of *N. benthamiana* (NbLab360.genome.fasta.gz), *S. lycopersicum* (S_lycopersicum_chromosomes.4.00.fa) and *S. aethiopicum* (Saethiopicum.fasta) were retrieved from public databases. Available gene annotations of *N. benthamiana* (NbLab360v103, Niben261 and Niben101), and the tomato (ITAG4.1) were used. To confirm gene expression, NCBI SRA transcriptomic datasets of *N. benthamiana* (SRR11282461, SRR11282462, SRR11282463 SRR24114147), *S. lycopersicum* (SRR28956637) and *S. aethiopicum* (SRR2229192, SRR6431645) were obtained from the Amazon Web Services (AWS) cloud using kingfisher. Adapter sequences and low-quality sequences were filtered using fastp to yield clean reads, which were mapped to the reference genome using the spliced aligner HISAT2 with the `--dta` option; obtained alignment .bam files were visualized using IGV and per base coverage of genomic loci were obtained using samtools `depth`. *De novo* transcriptome assembly was done using rnaSPAdes, a pipeline implemented within the SPAdes package. Transcripts of interest from assembled transcriptomes and available gene annotations were identified by DIAMOND `blastx` searches against a custom protein database.

Multi-species transcripts and genomic loci were aligned using MAFFT to identify conserved regions for vsRNAi design; transcript sequences used for vsRNAi design are shown in Data S11.

Dated molecular phylogeny for the Solanaceae was reported (Messeder *et al.*, 2024).

#### Viral vector systems

The tobacco rattle virus (TRV) vector system JoinTRV was used, which relies on pLX-TRV1 (Addgene:180515) and pLX-TRV2 (Addgene:180516), two T-DNA vectors of the pLX series (Pasin *et al.*, 2017) for simultaneous *Agrobacterium*-mediated inoculation of TRV genomic components (Aragonés *et al.*, 2022). A pLX-TRV2 derivative including an *N. benthamiana CHLI* cDNA fragment of 300 nt was described before (Aragonés *et al.*, 2022). An alternative TRV system consisting of pTRV1 (Addgene:148968) and pTRV2 (Addgene:148969) was also used (Liu *et al.*, 2002).

**One-step assembly of vsRNAi vectors**

For one-step assembly of vsRNAi into pLX-TRV2, oligonucleotide pairs were synthesized as follows: (i) a first oligonucleotide consisting, from the 5' end, of the TGAA sequence followed by the target gene sequence in the antisense orientation, and (i) a second oligonucleotide consisting, from the 5' end, of AGAG followed by the target gene sequence in the sense orientation. Custom DNA oligonucleotides were purchased desalted after synthesis as 100 mM solutions (Integrated DNA Technologies; Table S3). To recover oligonucleotide duplexes with overhangs compatible with BsaI-digested pLX-TRV2 (Table S4), pairs were mixed (10 µM each) in 1× T4 DNA ligase buffer (New England Biolabs) in a final volume of 10 µL, then incubated at 95°C (5 min), and cooled to 25°C (−0.1°C/s) and diluted by adding 190 µL of ultrapure water.

Recovered duplexes were assembled into pLX-TRV2 by one-step digestion-ligation reactions (Marillonnet and Grützner, 2020) including 1 µL duplex, ≈50 ng of pLX-TRV2, 0.5 µL T4 DNA ligase (M0202S, New England Biolabs), 0.5 µL BsaI-HFv2 (R3733S, New England Biolabs), 1× T4 DNA ligase buffer (New England Biolabs) in a final volume of 10 µL. Reaction mixtures were subjected to 30 cycles (5 min at 37°C and 5 min at 16°C), followed by 5 min at 60°C. Reactions were then taken to ice and transformed in *E. coli* DH5α competent cells, which were spread onto medium plates supplemented with 50 mg/L kanamycin and 40 µL 5-bromo-4-chloro-3-indolyl β-D-galactopyranoside (X-Gal; 40 mg/mL stock), and incubated (37°C, ≈24 h). White colonies were selected and recovered DNA plasmids were verified by Sanger sequencing.

For one-step assembly of vsRNAi into pTRV2, the duplex obtained using the oligonucleotide pair vCHLI-DK_F/vCHLI-DK_R (Tables S3, S4) was assembled into pTRV2 by a one-step digestion-ligation reaction in which BsaI-HFv2 was replaced with MluI (ER0561, Thermo Fisher Scientific); the resulting vector pTRV2-vCHLI with the *CHLI* sequence in the antisense orientation (vCHLI-DK) was identified by Sanger sequencing of recovered DNA plasmids.

The assembled vsRNAi vectors are summarized in Table S4, and their sequences are shown in Data S12.

**Viral vector delivery**

*Agrobacterium* AGL1 was used for viral vector delivery to plants (Pasin, 2022). In the JoinTRV system, AGL1 cells were electroporated with pLX-TRV1; colonies were selected on plates supplemented with rifampicin and gentamicin. Competent cells of the recovered strain AGL1(pLX-TRV1) were then electroporated with pLX-TRV2 and its derivatives. AGL1 transformants simultaneously hosting the two vectors of TRV genomic components were selected on plates supplemented with rifampicin, gentamicin, and kanamycin. In the pTRV1 + pTRV2 system, rifampicin and kanamycin were used to select AGL1 transformants hosting the vectors and their derivatives. Antibiotics were used at final concentrations of 30 mg/L gentamicin, 50 mg/L kanamycin, and 50 mg/L rifampicin.

Suspensions of the transformed bacteria were prepared in 10 mM 2-(*N*-morpholino)ethanesulfonic acid hydrate (MES; M8250, Sigma-Aldrich), 10 mM $MgCl_2$, 150 µM 3´,5´-dimethoxy-4-hydroxyacetophenon (acetosyringone; D134406, Sigma-Aldrich), pH 5.6, as described (Senthil-Kumar and Mysore, 2014; Uranga *et al.*, 2023). Bacterial suspensions were adjusted to an optical density at 600 nm ($OD_{600}$) of 0.5 and infiltrated in the abaxial side of one leaf of 3-week-old *N. benthamiana* plants using 1-mL needle-less syringes. For *S. lycopersicum* and *S. aethiopicum*, bacterial suspensions were adjusted to OD=2 and applied to fully expanded cotyledons of 10-day-old seedlings.

**Chlorophyll quantification**

Chlorophyll fluorometric quantification was done using the DUALEX® optical leaf clip meter (FORCE-A) (Cerovic *et al.*, 2012); data were obtained from upper uninoculated leaves and measurement was done with the adaxial leaf side facing the light sources, as previously reported (Villanueva *et al.*, 2023).

**Total RNA purification**

Samples from upper non-inoculated leaves were ground in liquid nitrogen in a mortar, and total RNA was extracted from ≈100 mg of plant material powder with the FavorPrep™ plant total RNA mini kit (FAPRK 001, Favorgen); during the preparation a mix of 5 U RNase-free DNase I (EN0521, Thermo Fisher Scientific), 100 U RiboLock RNase inhibitor (EO0381, Thermo Fisher Scientific), 1× DNase I reaction buffer with $MgCl_2$ in a final volume of 50 µL was added on column and incubated 10 min at 25°C to remove genomic DNA. RNA was eluted in RNase-free ultrapure water, quantified using a NanoDrop 1000 (Thermo Fisher Scientific), and stored at −80°C until further use.

**RT-qPCR analysis**

For RT-qPCR analysis, cDNA was synthesized using 1.2 µg of purified RNA and the iScript™ gDNA Clear cDNA Synthesis Kit (1725035, Bio-Rad Laboratories), following manufacturer instructions. The cDNA samples were used in qPCR reactions that included gene-pair-specific primers (Table S3) and 1× SsoAdvanced™ Universal SYBR® Green SuperMix (1725271, Bio-Rad Laboratories) and were run on a QuantStudio™ 3 Real-Time PCR system (Thermo Fisher Scientific). Expression was normalized using *PP2A* as a reference gene (Liu *et al.*, 2012), and fold changes relative to the control condition were calculated by the $\Delta\Delta^{CT}$ method (Pasin *et al.*, 2020).

**Transcriptome and small RNA sequencing and data analysis**

For each experimental condition, three biological replicates were used for transcriptomic analysis (RNA-seq; Table S5) and small RNA sequencing (sRNA-seq; Table S6); total RNA samples were processed at BGI (Hong Kong, China).

For RNA-seq, polyA-tailed RNA molecules were enriched and cDNA sequencing libraries were prepared following the DNBSEQ Eukaryotic mRNA library protocol, and sequenced (2 x 100-nt paired-end reads) on a DNBSEQ platform. Adapter sequences and low-quality sequences were filtered using fastp to yield clean mRNA reads. Transcripts were quantified by Salmon using the options `–keepDuplicates -k 31`, and a decoy-aware transcriptome index built using *N. benthamiana* annotated CDS (NbLab360.v103.gff3.CDS.fasta.gz) and, as a decoy, the complete genomic sequence (NbLab360.genome.fasta.gz). The obtained transcript-per-million matrix was used to assess transcriptome variation of the samples analyzed by principal component analysis. Differential expression analysis was calculated by edgeR using Salmon transcript read counts and trimmed mean of M values (TMM) normalization; quasi-likelihood F-test was applied to identify genes that are differentially expressed between sample groups, and adjusted *p* values (FDR) were computed by the Benjamini–Hochberg method. To inspect sequencing depth of *CHLI* genomic loci, clean reads were mapped using the spliced aligner HISAT2 with the `--dta` option against the *N. benthamiana* genome (NbLab360.genome.fasta.gz); obtained alignment .bam files were visualized using IGV and per base coverage were obtained using samtools `depth`. For each RNA-seq sample, per base coverage of *CHLI* genomic loci were normalized against the total number of mapped reads obtained by samtools `idxstats`.

For sRNA-seq, small RNA molecules were size enriched and cDNA sequencing libraries were prepared following the UMI small RNA library protocol, and sequenced (1 x 50-nt single-end read) on a DNBSEQ platform. Adapter sequences and low-quality sequences were filtered to yield clean sRNA reads. To analyze sRNA size classes, for each sample individual datasets with read length of 20, 21, 22, 23, 24, and 25 were obtained using fastp. Reads were mapped *to N. benthamiana* transcripts (NbLab360.v103.gff3.CDS.fasta.gz) using STAR, requiring at least an 18-nt perfect match (`--outFilterMatchNmin 18`) and 1 mismatch maximum (`--outFilterMismatchNmax 1`). Obtained alignment .bam files were visualized using IGV and per base coverage of transcripts were obtained using igvtools `count`. For each sRNA-seq sample, number of mapped reads per transcript and per base coverage of *CHLI* transcripts were normalized against the total number of mapped reads obtained by samtools `idxstats`, and reported as counts per million (CPM).

In plots showing the normalized sequencing depth per nucleotide, CPM values were obtained using the equation:

$$\frac{counts\ of\ reads\ mapped\ per\ nucleotide}{number\ of\ reads\ mapped\ to\ the\ reference\ dataset} \times 1\ M$$

wherein, the reference dataset consisted in the complete plant genome, transcriptome or the TRV RNA1 and RNA2 sequences from pLX-TRV1 and pLX-TRV2-vCHLI.

**Gene ontology term enrichment**

Gene ontology (GO) terms associated to *N. benthamiana* transcripts were retrieved by searching an *A. thaliana* protein database (Araport11_pep_20240409_representative_gene_model.gz) with DIAMOND `blastx` and retaining one target sequences per query (`-k 1`); GO annotation was then obtained from the Bioconductor database org.At.tair.db via topGO. Enrichment analysis of GO terms of biological processes in a transcriptome-wide analysis (global) was done by Kolmogorov-Smirnov testing considering edgeR computed FDR values; enrichment significance of GO terms of differentially expressed gene lists was determined by Fisher's exact test.

**Statistics**

Student’s *t*-test was used for pairwise comparisons; one-way ANOVA and Tukey’s honestly significant difference (HSD) test were used for multiple comparisons. Pearson correlation coefficient was calculated to estimate the linear relationship between the two variables. In differential expression analysis, the Benjamini–Hochberg multiple testing control method was used to compute adjusted *p* values (FDR); enrichment significance of GO terms was determined by Kolmogorov-Smirnov or Fisher's exact testing. Significance levels of *p* values are indicated in the figures and supporting information.

### SUPPORTING TABLES

**Table S1.** Transcriptomics and genomics resources used in the study.

| Dataset | Source | Identifier |
|---|---|---|
| *N. benthamiana* transcriptomic and small RNA sequencing reads for vsRNAi and control samples | This study | NCBI BioProject PRJNA1217923 |
| *N. benthamiana* genome and annotations | (Ranawaka *et al*., 2023) | Assembly: https://solgenomics.net/ftp/genomes/Nicotiana_benthamiana/LAB360/NbLab360.genome.fasta.gz<br><br>NbLab360v103: https://solgenomics.net/ftp/genomes/Nicotiana_benthamiana/LAB360/NbLab360.v103.gff3.CDS.fasta.gz<br><br>Niben261: https://solgenomics.net/ftp/genomes/Nicotiana_benthamianaV261/Nbenthamiana_Annotation/Niben261_genome.annotation.transcripts.fasta.gz<br><br>Niben101: https://solgenomics.net/ftp/genomes/Nicotiana_benthamiana/annotation/Niben101/Niben101_annotation.transcripts.fasta.gz |
| *N. benthamiana* transcriptomic datasets | (Pasin *et al*., 2020; Yue *et al*., 2023) | NCBI SRA: SRR11282461 SRR11282462 SRR11282463 SRR24114145 SRR24114146 SRR24114147 |
| *N. benthamiana* VIGS PDS transcriptomic datasets | (Ahmed *et al*., 2020) | NCBI SRA: SRR8691569 SRR8691570 SRR8691571 SRR8691565 SRR8691572 SRR8691574 |
| *N. benthamiana* VIGS RPL10 transcriptomic datasets | (Ahmed *et al*., 2020) | NCBI SRA: SRR8997267 SRR8997268 SRR8997269 SRR8997270 |
| *N. benthamiana* VIGS NRCX transcriptomic datasets | (Adachi *et al*., 2023) | NCBI SRA: ERR10087757 ERR10087758 ERR10087759 ERR10087760 ERR10087761 ERR10087762 |
| *S. lycopersicum* genome and annotations | (Hosmani *et al*., 2019) | Assembly: https://solgenomics.net/ftp/genomes/Solanum_lycopersicum/assembly/current_build/S_lycopersicum_chromosomes.4.00.fa<br><br>ITAG4.1 https://solgenomics.net/ftp/genomes/Solanum_lycopersicum/Heinz1706/annotation/ITAG4.1_release/ITAG4.1_cDNA.fasta |
| *S. lycopersicum* transcriptomic dataset | (Giovannini *et al*., 2024) | NCBI SRA: SRR28956637 |
| *S. aethiopicum* genome | (Benoit *et al*., 2025) | https://cshl-lippmanlab.s3.amazonaws.com/SolPanGenomics/Saethiopicum.genome_annotation.gz |
| *S. aethiopicum* transcriptomic datasets | (Gramazio *et al*., 2016; Wei *et al*., 2020) | NCBI SRA: SRR2229192 SRR6431645 |

| | | |
|---|---|---|
| *Arabidopsis thaliana* proteome | (Cheng *et al.*, 2017) | Araport11 https://www.arabidopsis.org/download/file?path=Sequences/Araport11_blastsets/Araport11_pep_20240409_representative_gene_model.gz |

**Table S2.** Software and algorithms used in the study.

| Software/Algorithm | Source | Identifier |
|---|---|---|
| DIAMOND | (Buchfink *et al.*, 2021) | https://github.com/bbuchfink/diamond |
| kingfisher | (Woodcroft *et al.*, 2024) | https://github.com/wwood/kingfisher-download |
| fastp | (Chen *et al.*, 2018) | https://github.com/OpenGene/fastp |
| HISAT2 | (Kim *et al.*, 2019) | http://daehwankimlab.github.io/hisat2/ |
| SPAdes | (Bushmanova *et al.*, 2019) | https://github.com/ablab/spades |
| MAFFT | (Katoh *et al.*, 2019) | https://mafft.cbrc.jp/ |
| salmon | (Patro *et al.*, 2017) | https://github.com/COMBINE-lab/salmon |
| STAR | (Dobin *et al.*, 2013) | https://github.com/alexdobin/STAR |
| samtools | (Li *et al.*, 2009) | http://www.htslib.org/ |
| edgeR | (Robinson *et al.*, 2010) | https://www.bioconductor.org/packages/edgeR |
| topGO | N/A | https://doi.org/doi:10.18129/B9.bioc.topGO |
| Integrative Genomics Viewer (IGV) | (Thorvaldsdóttir *et al.*, 2013) | https://igv.org/ |

**Table S3.** Oligonucleotides used in the study.

| ID | Sequence[a] | Use |
|---|---|---|
| vCHLI_F | TGAAACTTGGGCATGCATTCCAAATCGATCAAGAAG | Cloning |
| vCHLI_R | AGAGCTTCTTGATCGATTTGGAATGCATGCCCAAGT | Cloning |
| vCHLI_20_F | TGAAGCATGCATTCCAAATCGATC | Cloning |
| vCHLI_20_R | AGAGGATCGATTTGGAATGCATGC | Cloning |
| vCHLI_24_F | TGAAGGGCATGCATTCCAAATCGATCAA | Cloning |
| vCHLI_24_R | AGAGTTGATCGATTTGGAATGCATGCCC | Cloning |
| vCHLI_28_F | TGAATTGGGCATGCATTCCAAATCGATCAAGA | Cloning |
| vCHLI_28_R | AGAGTCTTGATCGATTTGGAATGCATGCCCAA | Cloning |
| vCHLI-b_F | TGAACTCATTACTTCTTGGTCATCTGGATCTGAATT | Cloning |
| vCHLI-b_R | AGAGAATTCAGATCCAGATGACCAAGAAGTAATGAG | Cloning |
| vPDS_F | TGAACAACATAGACTGATTGGGGTTGTAATATTCCT | Cloning |
| vPDS_R | AGAGAGGAATATTACAACCCCAATCAGTCTATGTTG | Cloning |
| vCHLI-DK_F | CGCGATTGAAACTTGGGCATGCATTCCAAATCGATCAAGAAGCTCT | Cloning |
| vCHLI-DK_R | CGCGAGAGCTTCTTGATCGATTTGGAATGCATGCCCAAGTTTCAAT | Cloning |
| CHLI_F | GGAGGAAGTTTTATGGAGGGATTAG | RT-qPCR |
| CHLI_R | GGATCAATTACATTCAGCAAAAGACA | RT-qPCR |
| PP2A_F | TGGGGATGGCTCCTGTTTTG | RT-qPCR |
| PP2A_R | CTCGCCAATGCCTGTCCTCT | RT-qPCR |

[a] Sequences are in the 5’ to 3’ orientation; overhangs compatible with BsaI-digested pLX-TRV2 are underlined

**Table S4.** Oligonucleotide duplexes used for vsRNAi vector assembly.

<table>
<tr><th>Oligonucleotide pair</th><th>Sequence</th><th>Assembled vector</th><th>Vector code</th></tr>
<tr><td>vCHLI_F<br><br>vCHLI_R</td><td>5' TGAAACTTGGGCATGCATTCCAAATCGATCAAGAAG 3'<br>||||||||||||||||||||||||||||||||<br>3' TGAACCCGTACGTAAGGTTTAGCTAGTTCTTCGAGA 5'</td><td>pLX-TRV2-vCHLI</td><td>vCHLI</td></tr>
<tr><td>vCHLI_20_F<br><br>vCHLI_20_R</td><td>5' TGAAGCATGCATTCCAAATCGATC 3'<br>||||||||||||||||||||<br>3' CGTACGTAAGGTTTAGCTAGGAGA 5</td><td>pLX-TRV2-vCHLI-20</td><td>vCHLI-20</td></tr>
<tr><td>vCHLI_24_F<br><br>vCHLI_24_R</td><td>5' TGAAGGGCATGCATTCCAAATCGATCAA 3'<br>||||||||||||||||||||||||<br>3' CCCGTACGTAAGGTTTAGCTAGTTGAGA 5'</td><td>pLX-TRV2-vCHLI-24</td><td>vCHLI-24</td></tr>
<tr><td>vCHLI_28_F<br><br>vCHLI_28_R</td><td>5' TGAATTGGGCATGCATTCCAAATCGATCAAGA 3'<br>||||||||||||||||||||||||||||<br>3' AACCCGTACGTAAGGTTTAGCTAGTTCTGAGA 5'</td><td>pLX-TRV2-vCHLI-28</td><td>vCHLI-28</td></tr>
<tr><td>vCHLI-b_F<br><br>vCHLI-b_R</td><td>5' TGAACTCATTACTTCTTGGTCATCTGGATCTGAATT 3'<br>||||||||||||||||||||||||||||||||<br>3' GAGTAATGAAGAACCAGTAGACCTAGACTTAAGAGA 5'</td><td>pLX-TRV2-vCHLI-b</td><td>vCHLI-b</td></tr>
<tr><td>vPDS_F<br><br>vPDS_R</td><td>5' TGAACAACATAGACTGATTGGGGTTGTAATATTCCT 3'<br>||||||||||||||||||||||||||||||||<br>3' GTTGTATCTGACTAACCCCAACATTATAAGGAGAGA 5'</td><td>pLX-TRV2-vPDS</td><td>vPDS</td></tr>
<tr><td>vCHLI-DK_F<br><br>vCHLI-DK_R</td><td>5' CGCGATTGAAACTTGGGCATGCATTCCAAATCGATCAAGAAGCTCT 3'<br>||||||||||||||||||||||||||||||||||||||||||<br>3' TAACTTTGAACCCGTACGTAAGGTTTAGCTAGTTCTTCGAGAGCGC 5'</td><td>pTRV2-vCHLI</td><td>vCHLI-DK</td></tr>
</table>

175

**Table S5.** Summary of the transcriptome sequencing (RNA-seq) datasets generated.

| Condition | Biological replicate | Clean read pairs (2x 100) | Clean base | Sample description |
|---|---|---|---|---|
| CTRL | 1 | 34643286 | 6.842E+09 | Total RNA sample from upper uninoculated leaves of *Nicotiana benthamiana* plants inoculated with pLX-TRV1 and the pLX-TRV2 empty vector, and collected after 13 days; polyA-tailed RNA molecules were enriched and sequenced (PE100). |
| CTRL | 2 | 31841406 | 6.295E+09 | |
| CTRL | 3 | 34620206 | 6.835E+09 | |
| vCHLI | 1 | 34650755 | 6.843E+09 | Total RNA sample from upper uninoculated leaves of *Nicotiana benthamiana* plants inoculated with pLX-TRV1 and pLX-TRV2-vCHLI, and collected after 13 days; polyA-tailed RNA molecules were enriched and sequenced (PE100). |
| vCHLI | 2 | 34583130 | 6.826E+09 | |
| vCHLI | 3 | 34620846 | 6.835E+09 | |

**Table S6.** Summary of the small RNA sequencing (sRNA-seq) datasets generated.

| Condition | Biological replicate | Clean read count | Average read length (nt) | Sample description |
|---|---|---|---|---|
| CTRL | 1 | 12153131 | 19.67 | Total RNA sample from upper uninoculated leaves of *Nicotiana benthamiana* plants inoculated with pLX-TRV1 and the pLX-TRV2 empty vector, and collected after 13 days; small RNA molecules were size enriched and sequenced (SE50). |
| CTRL | 2 | 9775190 | 19.85 | |
| CTRL | 3 | 11982178 | 19.21 | |
| vCHLI | 1 | 10562799 | 19.44 | Total RNA sample from upper uninoculated leaves of *Nicotiana benthamiana* plants inoculated with pLX-TRV1 and pLX-TRV2-vCHLI, and collected after 13 days; small RNA molecules were size enriched and sequenced (SE50). |
| vCHLI | 2 | 10891736 | 19.88 | |
| vCHLI | 3 | 11735408 | 20.17 | |

### SUPPORTING FIGURES

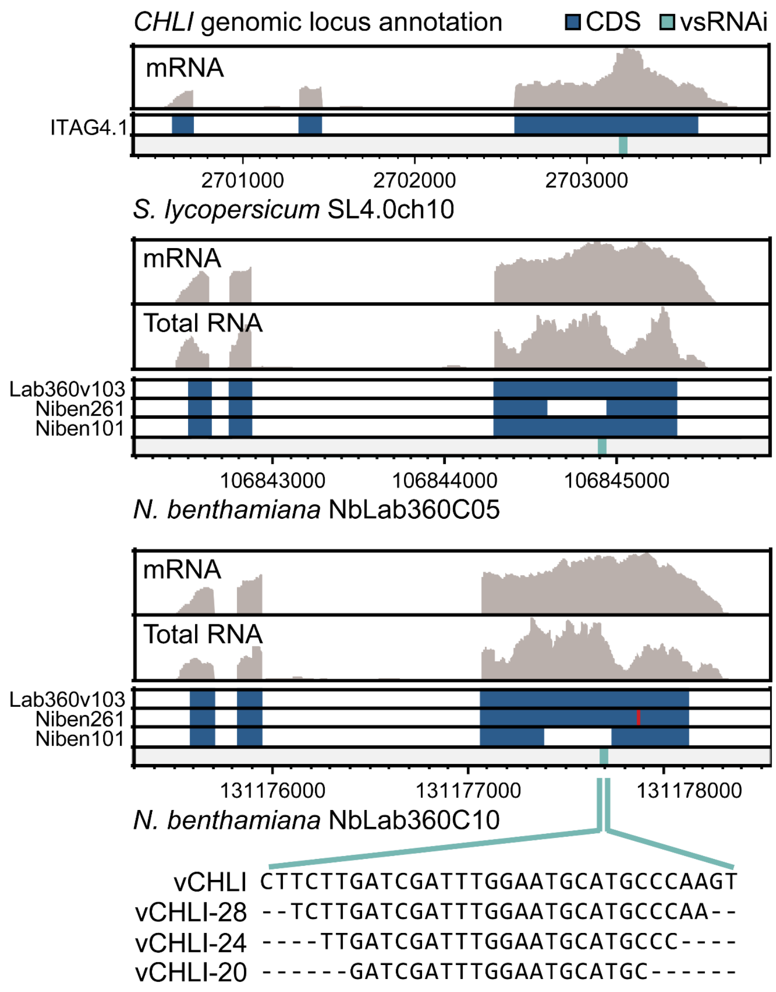


**Figure S1. Annotation of *Nicotiana benthamiana CHLI* genomic loci and design of vsRNAi.** Comparative genomics and transcriptomics guided the design of vsRNAi of 32, 28, 24 and 20 nt (vCHLI, vCHLI-28, vCHLI-24, and vCHLI-20) with sequences conserved in tomato (*S. lycopersicum*) and *N. benthamiana CHLI* genomic loci. Chromosomal regions were identified by DIAMOND blastx searches; for each genomic locus, expression and gene structure were confirmed by mapping sequencing reads obtained by either rRNA-depleted total RNA (Total RNA) or polyA-tailed RNA (mRNA) samples. Coding sequences (CDS) from previously reported annotations are shown as blue boxes; the red line indicates a sequence insertion unsupported by transcriptomic analysis.

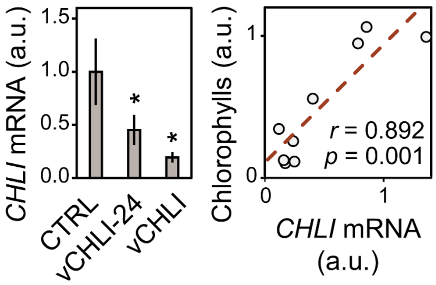


**Figure S2. RT-qPCR of *N. benthamiana CHLI* transcripts.**

Left, *N. benthamiana CHLI* transcript quantification in vCHLI, vCHLI-24 and control (CTRL) samples (mean ± SD, $n$ = 3); significance levels versus the control condition as per Student's *t*-test are shown; ∗, $p < 0.05$. Right, linear relationship between *CHLI* transcript and chlorophyll amounts ($n$ = 9); Person's $r$ and $p$ value are indicated.

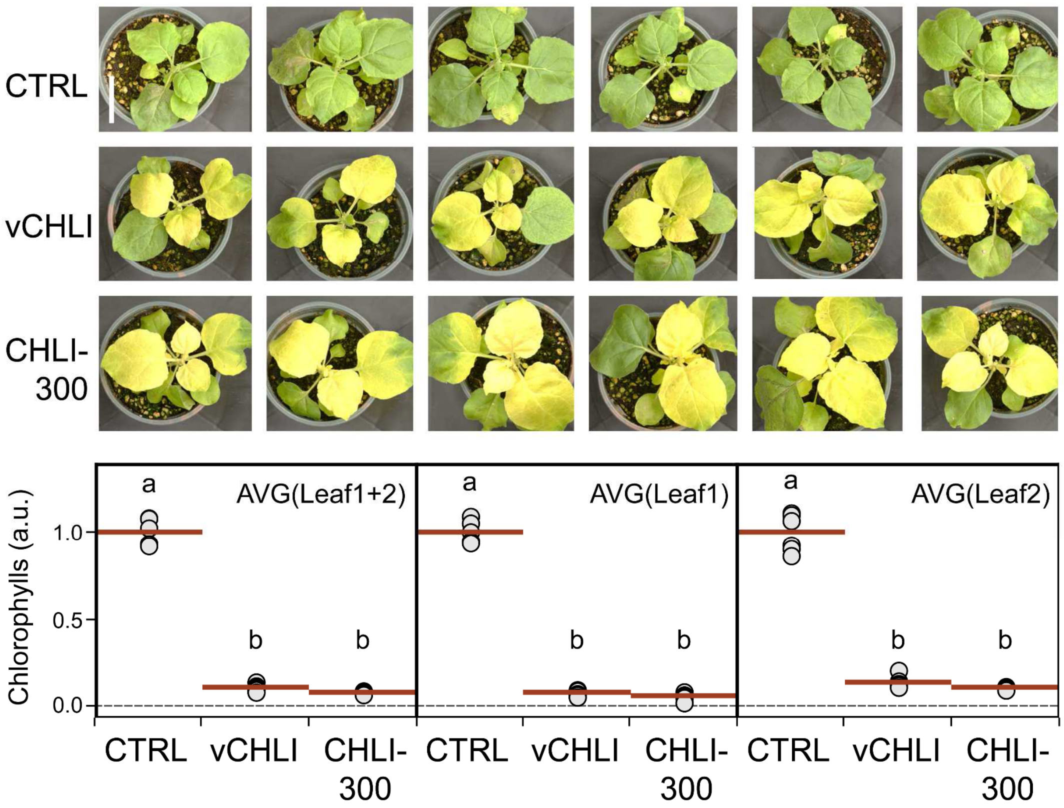


**Figure S3. Delivery of a 32-nt vsRNAi triggers robust silencing phenotypes.**
Plants were treated with the unmodified JoinTRV (CTRL), or derivatives including a 32-nt vsRNAi (vCHLI) targeting the two *N. benthamiana CHLI* homeologues, and a 300-nt *CHLI* cDNA fragment previously reported (Aragonés *et al.*, 2022). Top, plant phenotypes (scale = 5 cm). Bottom, chlorophyll levels, with each condition represented by a red bar (average of 6 plants). Individual dots show the average chlorophyll content of a single plant, measured at the most and second-most symptomatic leaf, and both leaves combined (AVG(Leaf1), AVG(Leaf2), and AVG(Leaf1+2), respectively). For control plants, equivalent leaves were measured. Five measurements were taken per leaf and averaged to calculate the values shown. Different letters indicate significant differences ($p < 0.05$), by one-way ANOVA and Tukey's HSD test.

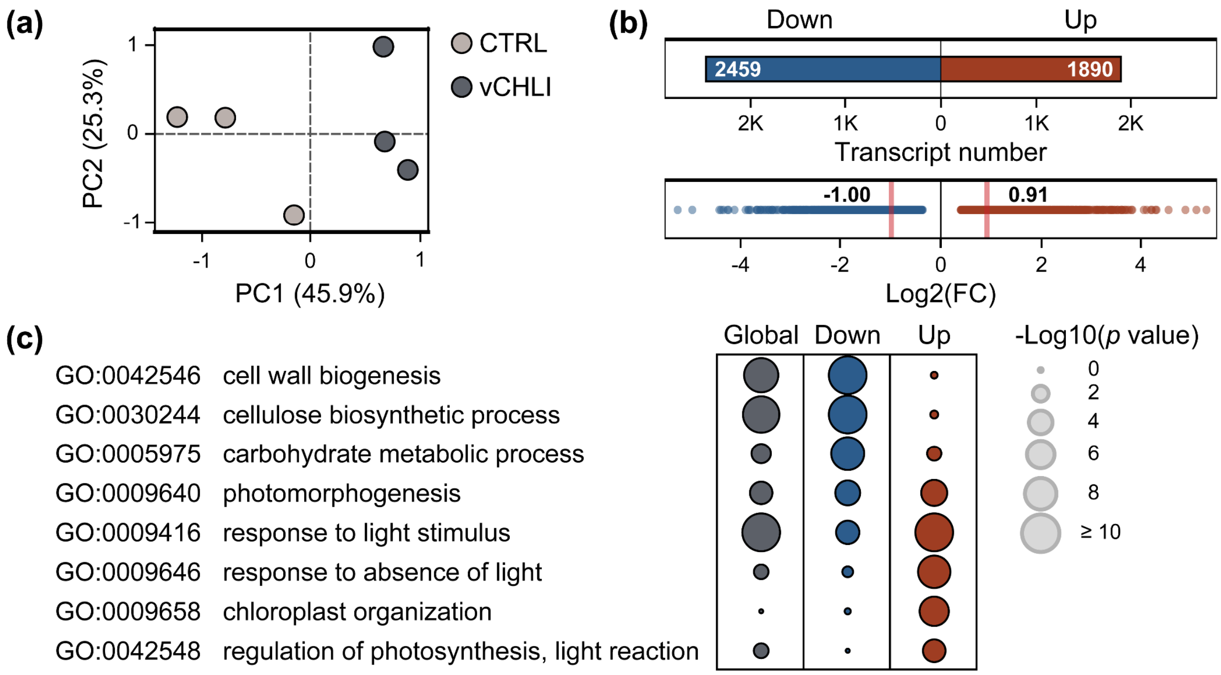


**Figure S4. Functional analysis of transcriptome-wide changes triggered by vsRNAi.**

(**a**) Principal component analysis of samples analyzed by transcriptomics; CTRL, control condition samples; vCHLI, samples from *N. benthamiana* plants inoculated with TRV vector comprising a 32-nt vsRNAi targeting *CHLI* homeologues ($n = 3$).

(**b**) Total number (top) and fold change (bottom; median values are shown) of differentially expressed transcripts (FDR < 0.05; Data S1, Data S2).

(**c**) Enrichment of gene ontology (GO) terms of biological processes in a transcriptome-wide analysis (global; Data S3), and in subset lists including down- or up-regulated transcripts (Data S4, S5); representative terms are shown with their significance levels.

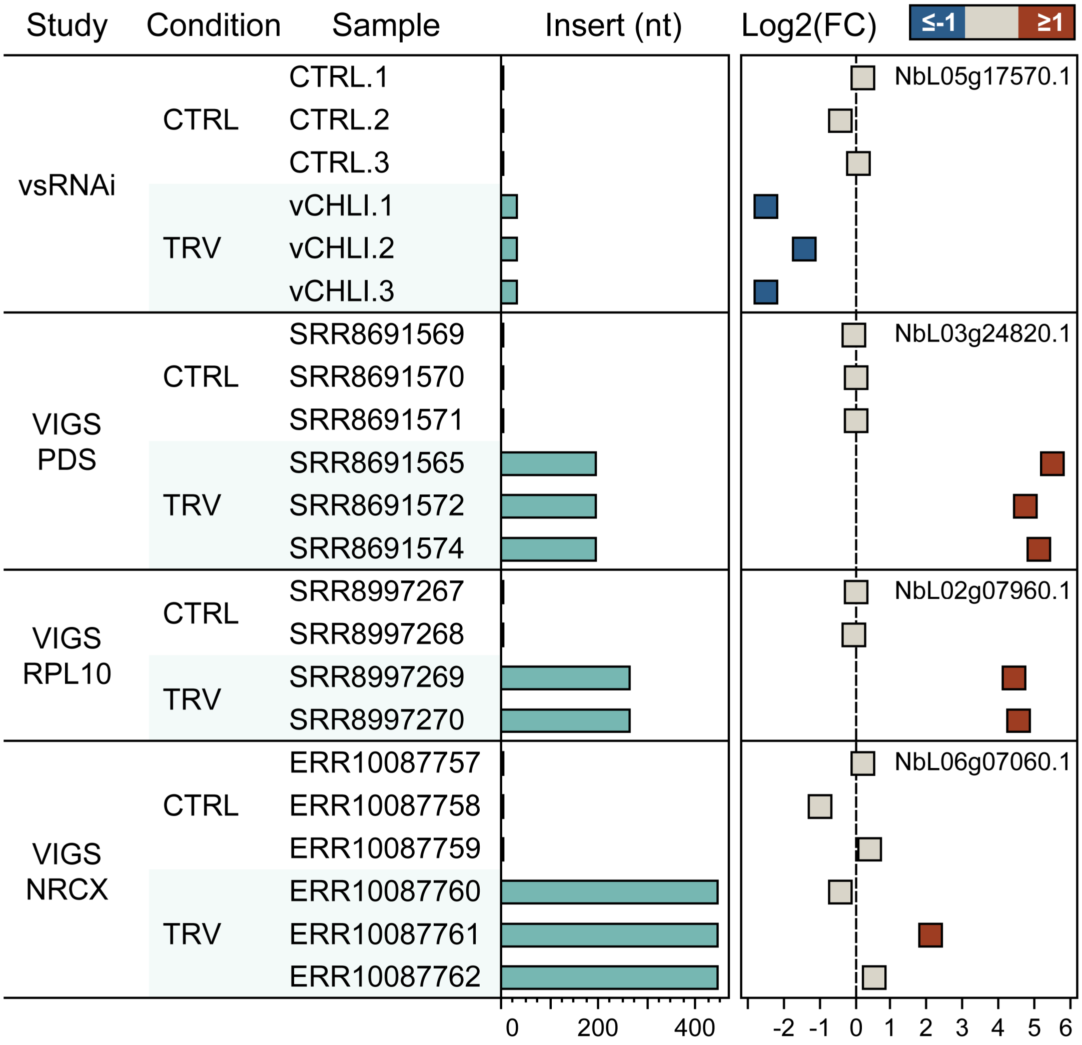


**Figure S5. vsRNAi enable transcriptome-wide quantification of target gene silencing.**
Transcriptomes of *N. benthamiana* plants treated with TRV vectors containing vsRNAi or larger VIGS inserts were analyzed using the alignment-free method Salmon (Patro *et al.*, 2017). The obtained transcript-per-million (TPM) values were used to calculate the abundance of target genes compared to control samples, and expressed as log2 fold change (FC). Target gene downregulation was confirmed in vsRNAi samples from this study, whereas no downregulation was detected in the VIGS studies analyzed (PDS, RLP10, and NRCX). VIGS transcriptomes and the insert sizes were reported (Adachi *et al.*, 2023; Ahmed *et al.*, 2020); NCBI SRA identifiers of the RNA-seq samples analyzed are shown.

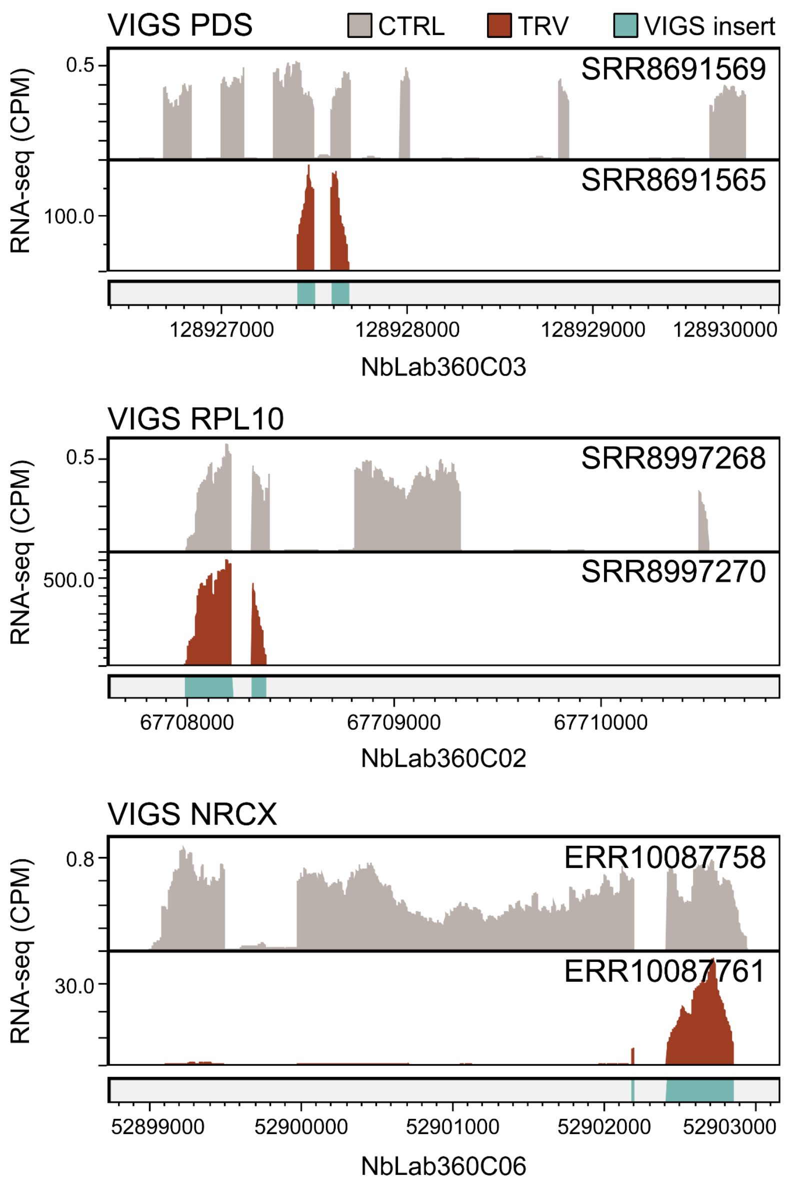


**Figure S6. Viral amplification of large VIGS inserts interferes with quantification of target genes.**

RNA-seq reads from TRV VIGS (TRV) and control (CTRL) samples were mapped to the *N. benthamiana* genome using the HISAT2 aligner (Kim *et al.*, 2019). Plots show the number of reads, expressed as counts per million (CPM), mapped to the genomic loci of target genes. TRV samples showed high CPM values within the genomic regions that share homology with the VIGS inserts, which greatly exceeded those of control samples. VIGS transcriptomes were reported (Adachi *et al.*, 2023; Ahmed *et al.*, 2020), and NCBI SRA identifiers of the datasets analyzed are shown within each plot.

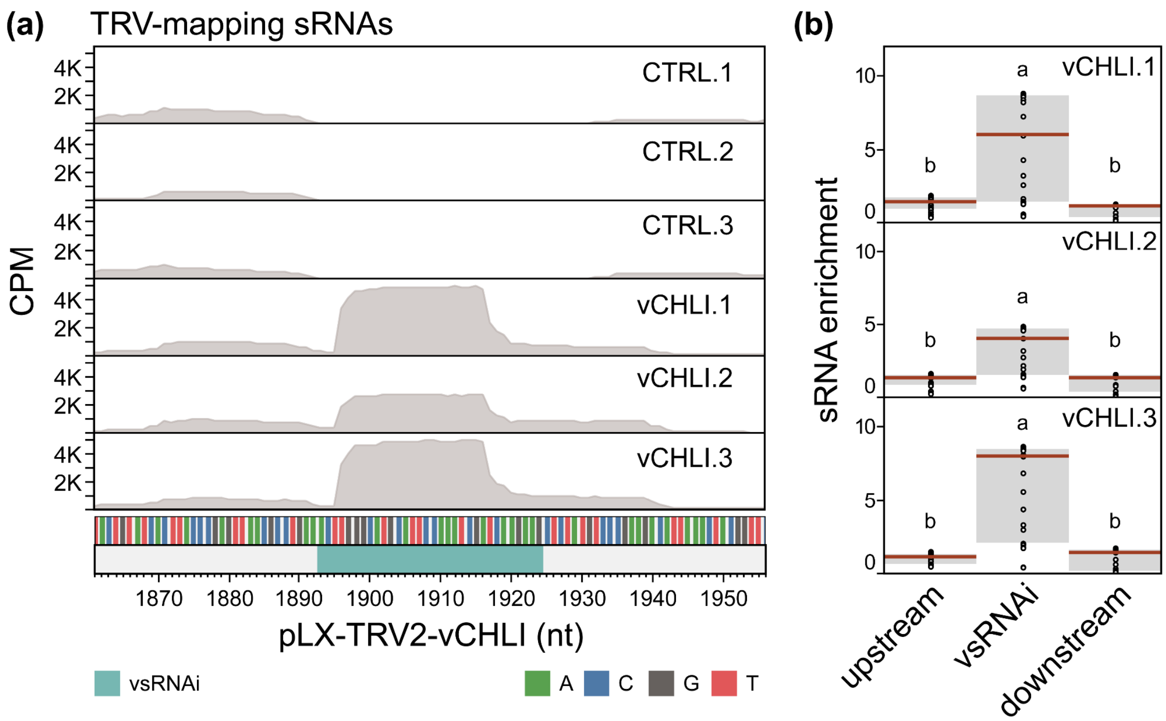


**Figure S7. vsRNAi trigger host-derived production of sRNAs.**

(**a**) Small RNA sequencing (sRNA-seq) reads of vCHLI and CTRL samples were mapped to the TRV RNA2 sequence of pLX-TRV2-vCHLI, including a 32-nt vsRNAi targeting *CHLI* homeologues. Plots show the number of mapped reads. vCHLI samples showed high CPM values spanning the vsRNAi position, which greatly exceeded those of control samples.

(**b**) Enrichment of sRNA mapping to vsRNAi compared to flanking viral sequences. The 32-nt vsRNAi sequence had significantly higher per nucleotide CPM values than the 32-nt viral sequences immediately up- and downstream, indicating viral-independent and host-derived sRNA production. Letters indicate $p < 0.05$, one-way ANOVA and Tukey's HSD test.

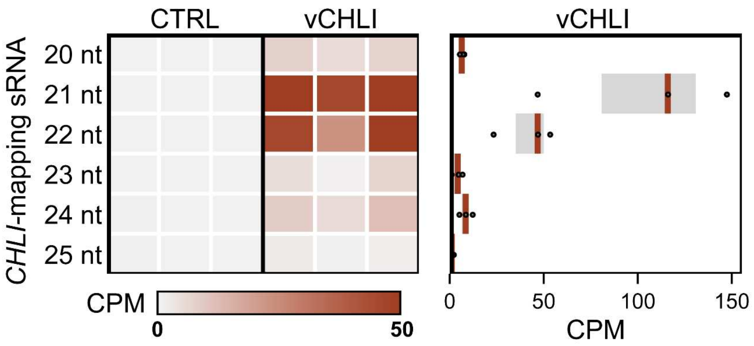


**Figure S8. Production of gene-specific sRNAs triggered by vsRNAi.**

Small RNA sequencing (sRNA-seq) reads of vCHLI and CTRL samples (*n* = 3) were mapped to *N. benthamiana CHLI* transcripts (Data S7, S8). Left, heatmap of counts per million (CPM) per sample and size class; right, plot of CPM values of vCHLI samples (median, upper and lower quartiles are marked; *n* = 3).

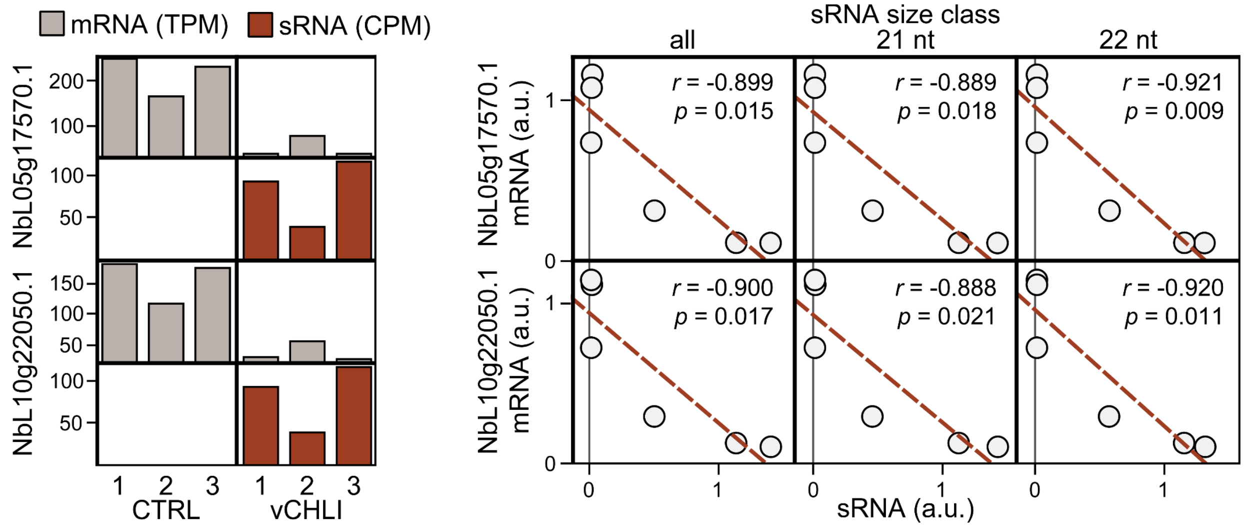


**Figure S9. Abundance of *CHLI*-mapping sRNAs negatively correlated with *CHLI* expression.**

Left, abundance of *CHLI* mRNA (transcript per million, TPM) and *CHLI*-mapping sRNAs (counts per million, CPM) in CTRL and vCHLI samples are shown for the two *CHLI* homeologues (NbL05g17570.1, NbL10g22050.1). Right, scatter plots show correlation ($n$ = 6) among the abundance of *CHLI*-mapping sRNAs and *CHLI* transcripts; Pearson correlation coefficients ($r$) and significance levels ($p$) are shown.

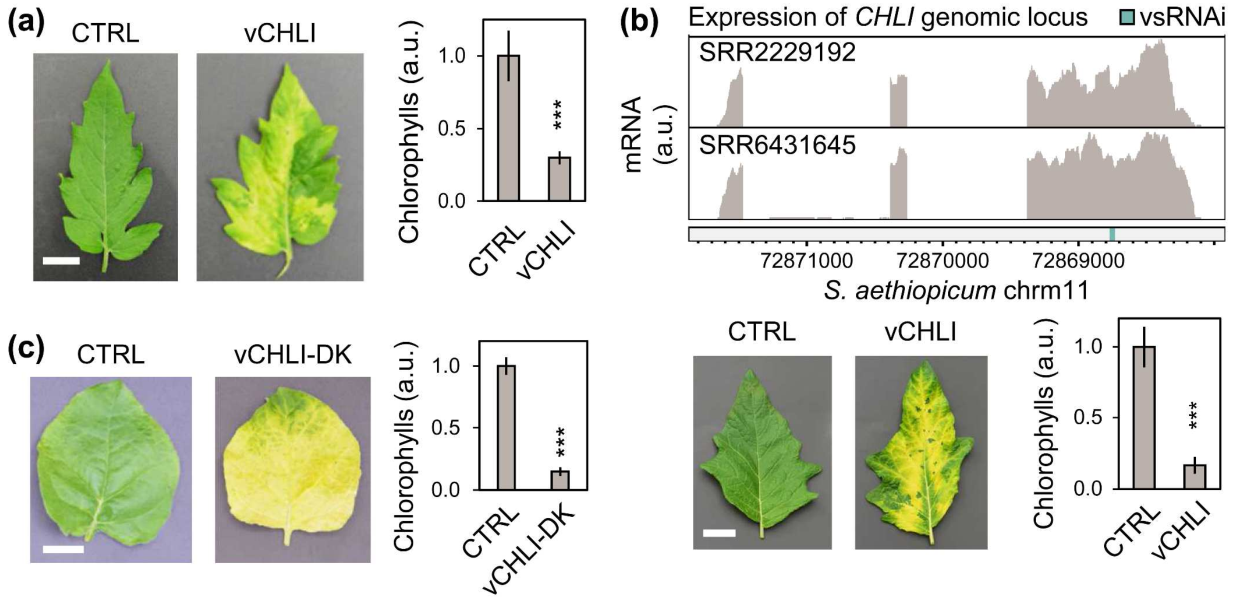


**Figure S10. Portability of vsRNAi approaches to crops and viral vector systems.**

**(a)** The JoinTRV derivative vCHLI causes a chlorophyll reduction in tomato (*S. lycopersicum*). Phenotypes of upper uninoculated leaflets of "Moneymaker" plants and chlorophyll fluorometric quantification results are shown (mean ± SD, $n = 4$; ∗∗∗, Student's *t*-test $p < 0.001$); CTRL, control; scale = 1 cm.

**(b)** The JoinTRV derivative vCHLI causes a chlorophyll reduction in scarlet eggplant (*Solanum aethiopicum*). Top, identification and annotation of the *S. aethiopicum CHLI* genomic locus; results of transcriptomic analysis and the vsRNAi targeted sequence of the identified *CHLI* genomic locus are shown. Bottom, phenotypes of upper uninoculated leaves of "Rossa di Rotonda" plants, and chlorophyll fluorometric quantification results are shown (mean ± SD, $n = 3$; ∗∗∗, Student's *t*-test $p < 0.001$); scale = 2.5 cm.

**(c)** Use of a 32-nt vsRNAi derivative of the pTRV1 + pTRV2 vector system from Liu *et al.* (2002) to target the *N. benthamiana CHLI* gene pairs (vCHLI-DK) results in a chlorophyll reduction. Phenotypes of *N. benthamiana* upper uninoculated leaves and chlorophyll fluorometric quantification results are shown (mean ± SD, $n = 3$; ∗∗∗, Student's *t*-test $p < 0.001$); scale = 2.5 cm.
